# Spatial quantitative metabolomics enables identification of remote and sustained ipsilateral cortical metabolic reprogramming after stroke

**DOI:** 10.1101/2024.10.26.620403

**Authors:** Gangqi Wang, Bernard M. van den Berg, Sarantos Kostidis, Kelsey Pinkham, Marleen E. Jacobs, Arthur Liesz, Martin Giera, Ton J. Rabelink

## Abstract

Mass spectrometry imaging (MSI) has become a cornerstone of spatial biology research. However, various factors that are intrinsic to the technology limit quantitative capacity of MSI-based spatial metabolomics and thus reliable interpretation. Here, we developed a quantitative MSI workflow (Q-MSI), based on isotopically ^13^C-labeled yeast extract as internal standards, to overcome these pitfalls. Using brain and kidney tissue, we demonstrate that this approach allows for (absolute) quantification of hundreds of metabolites and lipids. Applying our workflow to a stroke model allowed us to not only map metabolic remodeling of the infarct and peri-infarct area over time, but also discover hitherto unnoted remote metabolic remodeling in the histologically unaffected ipsilateral sensorimotor cortex. At day 7 post-stroke, increased levels of neuroprotective lysine and reduced excitatory glutamate levels were found compared to the contralateral cortex. By day 28 post-stroke, lysine and glutamate levels had been normalized, while decreased precursor pools of UDP-GlcNAc and linoleate persisted that have previously been associated with vulnerability. Importantly, traditional normalization strategies not employing internal standards were unable to visualize these differences. Using ^13^C-labeled yeast extracts as a normalization strategy establishes a new paradigm in quantitative MSI-based spatial metabolomics that greatly enhances reliability and interpretive strength.

## Main

Spatial biology has taken center stage in life science and biotechnology research. Spatially resolved molecular technologies are nowadays allowing researchers to shed light on the spatial heterogeneity of biological tissues, microenvironments, and cell-cell communications, integrating biology’s central disciplines, i.e. transcriptome, epigenome, proteome, and metabolome^1^. Spatial metabolomics, as an important component of spatial biology, provides insights into the in-situ distribution of metabolites and metabolic microenvironments. MSI based spatial metabolomics serves as the primary approach for examining the spatial distribution of metabolites^2^. However, several pitfalls as for example, ion suppression, adduct formation and in-source fragmentation render MSI based (absolute) quantification cumbersome if not impossible^3, 4^. These challenges are intrinsic to mass spectrometry (MS) based approaches and consequently affect both major MSI technologies, matrix assisted laser desorption ionization (MALDI) and desorption electrospray ionization (DESI)^5, 6^. Even more so, these limitations can affect both inter-tissue comparison and more critically regional comparisons within a single tissue^5^. Batch effects caused by day to day experimental and instrumental variations further complicate the matter^7^. In summary, (absolute) quantification and standardization of spatial metabolomics is jeopardized by various analytical intricacies that yet have to be overcome. The most widely accepted and practiced approach for enabling absolute quantification and reducing intra- and inter-batch variations is the use of isotopically labelled internal standards (IS), commonly using ^13^C labelled variants^3^. Such a strategy has recently been applied to spatial lipidomics analysis compensating for ion suppression and ensuring data reproducibility across different laboratories^8^. The method involves the use of class specific IS homogeneously applied to the tissue surface. The approach is facilitated by the fact that lipids follow a building block structure with lipid class chemical properties largely being dictated by the lipid headgroup^9^. Consequently, a limited number of only 13 class specific IS could serve as reference points for pixelwise normalization of each lipid species within the tissue sample. However, extrapolating the concept to spatial metabolomics studies presents challenges due to the more diverse physico-chemical properties of (water-soluble) metabolites compared to lipids. Even for distinct metabolite classes, physico-chemical properties and structures can vastly differ, necessitating metabolite rather than class specific IS. However, the use of extended metabolite IS panels can be costly or constrained by availability, particularly when working at an omics-scale. Alternatively, uniformly ^13^C-labeled yeast extracts offer a rich source of isotopically labeled metabolites derived from evolutionarily conserved primary metabolomes. These extracts have been used as IS for quantitative metabolomics using LC-MS and GC-MS^10, 11^. Relying on the biosynthetic machinery of yeast to generate a multitude of ^13^C labeled metabolite standards, we here introduce a quantitative spatial metabolomics method that combines ^13^C-labeled yeast extracts with MALDI-MSI, enabling pixelwise IS normalization and quantification of hundreds of metabolites and lipids.

This quantitative MSI method is based on previously validated “on-tissue” spraying approaches^12^. Uniformly ^13^C-labeled yeast extracts were homogeneously sprayed onto heat-inactivated tissue surface, followed by application of N-(1-naphthyl) ethylenediamine dihydrochloride (NEDC) matrix deposition as previously described^13^. Detection of in-situ metabolites in negative mode was performed using a TimsTOF flex MALDI2 mass spectrometer (Figure 1A). Tissues sprayed with and without ^13^C-labeled yeast extracts exhibited comparable spectrum quality, enabling the identification of yeast-derived ^13^C-labeled metabolites (Supplementary Figure 1). To investigate the overall ion suppression of metabolites, we performed molecular histology based on both yeast-derived ^13^C-labeled metabolites and endogenous lipids to visualize tissue structure in kidney and brain (Figure 1B). The exogenous metabolites exhibited histological signals similar to tissue structures (Figure 1B), while differently sprayed ^13^C-labeled metabolites displayed distinct ion suppression within each tissue sample (Supplementary Figure 2). These findings underscore the importance of correcting ion suppression for individual metabolites in spatial metabolomics studies. Overall, we identified 427 metabolite/lipid features present in both unlabeled and ^13^C-labeled yeast extracts, jointly serving as an IS library for subsequent quantification (Figure 1C). We examined the presence of both endogenous unlabeled metabolites and their corresponding exogenous ^13^C-labeled metabolites in kidney and brain tissues sprayed with ^13^C-labeled yeast extracts, excluding overlapping peaks. Overall, 171 and 170 metabolite/lipid features were found useful for pixelwise normalization on kidney and brain respectively, and 145 features could be applied to both tissues(Figure 1C). These metabolites are biochemically involved in glycolysis/gluconeogenesis, the TCA cycle, pentose phosphate pathway, amino acid and fatty acid metabolic pathways (Figures 1D). Specific examples include glutathione (GSH), glutathione disulfide (GSSG), Uridine diphosphate N-acetylglucosamine (UDP-GlcNAc) and reduced-nicotinamide adenine dinucleotide (NADH). In addition, the identified labelled IS also included more complex lipids which can serve to normalize individual lipid species (and classes). Subsequently we selected several lipid IS for class-wise normalization allowing the quantification of hundreds of lipid species belonging to the following lipid classes LPA, LPE, LPI, PE, PA, PG, PI and PS. For example, 146 lipid species from 8 lipid classes could be quantified on brain tissues. To test the IS we subsequently compared relative quantification with the commonly applied root mean square (RMS) and total ion count (TIC) normalization methods. While both traditional normalization methods demonstrated enhancements compared to no normalization, they presented with vastly different results when compared with individual ^13^C-labelled IS (Figures 1E-F).

**Figure 1.**
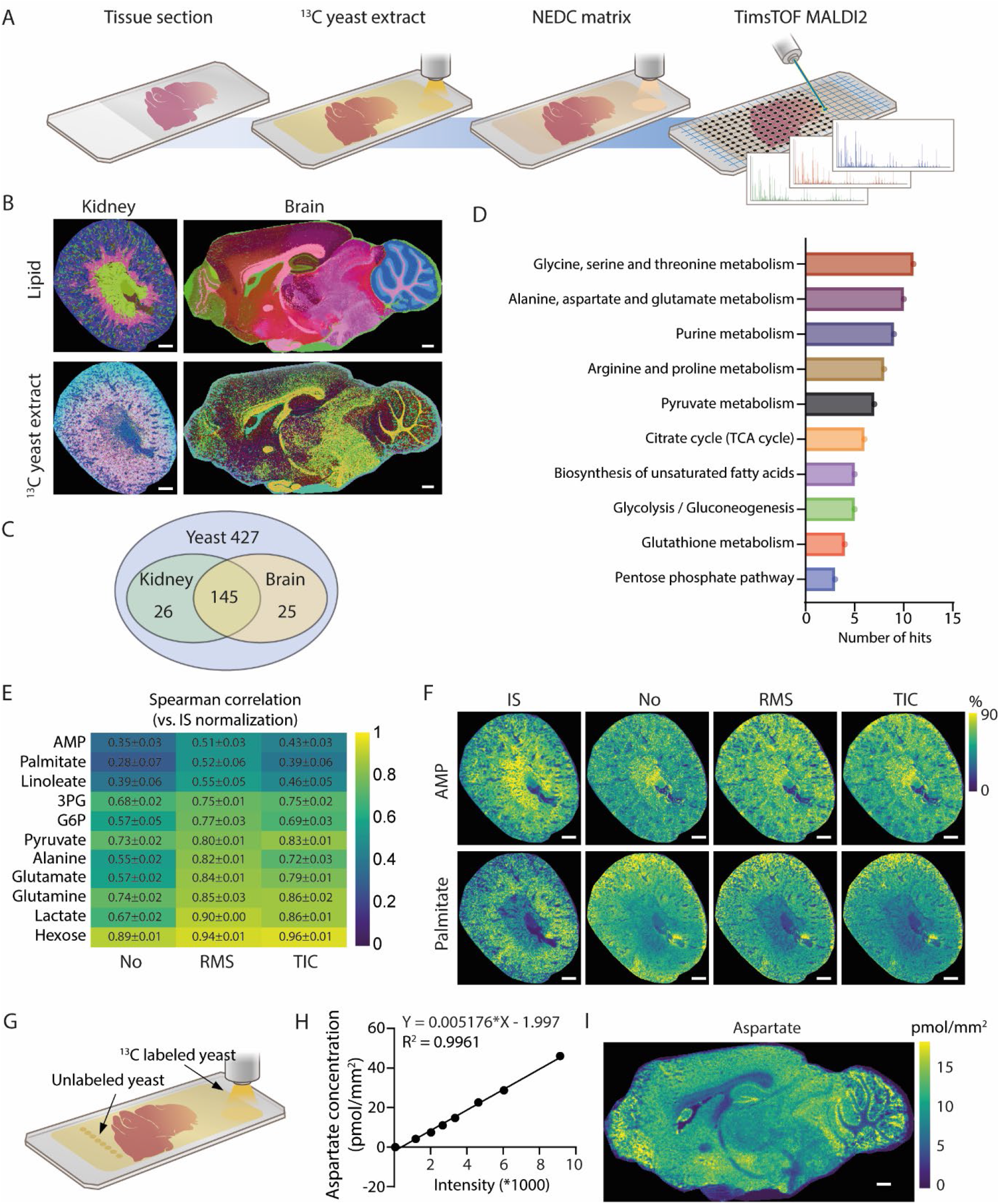
^13^C Yeast-assisted mass spectrometry imaging for quantitative spatial metabolomics. **A,** Workflow of quantitative spatial metabolomics.**B,** Comparison of molecular histology generated from endogenous lipids and exogenous ^13^C-labeled yeast extract. **C,** Amount of available internal standards from yeast extract and its availability on kidney and brain segments. **D,** KEGG pathway analysis on the quantitative metabolites of both kidney and brain. **E,** Spearman correlation analysis between internal standard (IS) normalization with root mean square (RMS), total ion count (TIC) or no normalization methods from kidney data. **F,** Distribution of AMP and palmitate using different normalization methods. **G,** Scheme of absolute quantitative spatial metabolomics. **H,** Calibration curve for aspartate quantification. **I,** Absolute spatial quantification of aspartate on mouse brain. All scale bars = 600 μm.

To investigate the method’s potential for absolute quantification, we initially determined metabolite concentrations in the unlabeled yeast extract using nuclear magnetic resonance spectroscopy. Subsequently, we spotted varying amounts of unlabeled yeast extracts on the slide alongside the tissue prior to spraying with ^13^C-labeled yeast extracts (Figure 1G). Based on this, a set of calibration curves were constructed for *in situ* quantification using the obtained response factors (Figure 1H and Supplementary Figure 3). This absolute quantification not only allows inter-region comparison of identical metabolites but also inter-metabolite comparison. Taking brain tissue as an example, the excitatory neurotransmitter aspartate showed a specific quantitative distribution in the cerebellar granular layer of approximately15 pmol/mm^2^ (Figure 1I), which is consistent with previous reports claiming that aspartate is a possible neurotransmitter in cerebellar climbing fibers^14^.

To demonstrate the potential of this quantitative method for biological discovery, we conducted region specific analysis on mouse brains in an experimental photothrombotic stroke model^15^. Samples from both day 7 post-stroke (subacute phase) and day 28 post-stroke (chronic phase) were analyzed. Integrating H&E staining on post-MSI tissue with spatial segmentation of the MALDI-MSI data, uniform manifold approximation and projection (UMAP) of lipid features revealed 5 distinct infarct (clusters 12, 21 and 26) and peri-infarct (clusters 20 and 25) regions (Figures 2A-B). In addition to metabolic heterogeneity in the infarct area, the peri-infarct and healthy regions, inter and intra-day 7 and day 28 post-stroke brains, revealed a metabolic gradient from the infarct core to the healthy tissue area (Figure 2A). Besides changes observed in the distribution of lipid species, a gradual increase in the excitatory neurotransmitter glutamate and decrease of GSSG/GSH ratio were observed from the necrotic core to healthy regions (Figures 2C-D), indicating altered neurological function and increased oxidative stress levels as a result of the insult. At day 28 post-stroke, the infarct region showed higher GSSG/GSH ratio and higher linoleate level compared to the healthy region, although GSSG/GSH ratio decreased below day 7 levels (Figures 2D-E). Linoleate could potentially further participate in regulating neurotransmission through its oxidized derivatives upon ischemic brain stroke^16^. Interestingly, in the absence of any histological changes we observed remote metabolic remodeling in the ipsilateral sensorimotor cortex region (cluster 16) of the stroke area, in comparison to the contralateral cortical region (cluster 6), (Figures 2A-B). This ipsilateral cortical region showed normal myelin coverage and cell number compared to its contralateral cortical region (Figures 2F), indicating an absence of structural brain damage. Based on morphology we could compare cluster 16 with its contralateral region located in the non-affected brain hemisphere within cluster 6 (Figure 2G). 23 out of 131 quantified metabolites and 61 out of 146 quantified lipids were significantly changed at day 7 post-stroke, and 37 metabolites and 59 lipids were significantly changed at day 28 post-stroke (Figure 2H). This distinct metabolic remodeling in the cluster 16 region at days 7 and 28 post-stroke showed different phases of metabolic remodeling (Figure 2H). Two distinct *m/z* values (436.283 and 464.314) abundantly present in cluster 16 over time (Figure 2I-J) were preliminarily annotated as LPE O-16:1 and LPE O-18:1 using database search and off-tissue LC-MS/MS analysis as described below. LPEs have been implicated in stimulating neurite outgrowth in cultured cortical neurons^17^, suggesting a potential contribution to stroke recovery in cluster 16 region. Decreased glutamate and increased lysine levels were observed in cluster 16 at day 7 post-stroke, which normalized at 28 days post stroke compared to control cluster 6 (Figures 2K-L). Glutamate has been shown to increase the affinity of acid-sensing ion channels for protons and is associated with aggravation of ischemic neurotoxicity^18^, while opposite lysine could inhibit glutamate-induced neuronal activity and is associated with reduction of the effects of cerebral ischemic insults^19^. These data suggest protective metabolic remodeling in the ipsilateral cortical region after hypoxic injury in the subacute phase of stroke. At day 28 post-stroke however, the O-GlcNAcylation precursor pools of UDP-GlcNAc and linoleate were significantly reduced (Figures 2M-N), suggesting long term sustained metabolic rewiring in the remote ipsilateral cortical region after stroke. UDP-GlcNAc availability is directly associated with O-GlcNAcylation of neuronal proteins and protection of ischemic injury^20^. Similarly, linoleate availability has been related to potential lower stroke risk^21^. Such longer term metabolic remodeling could render the brain vulnerable to further injury.

**Figure 2.**
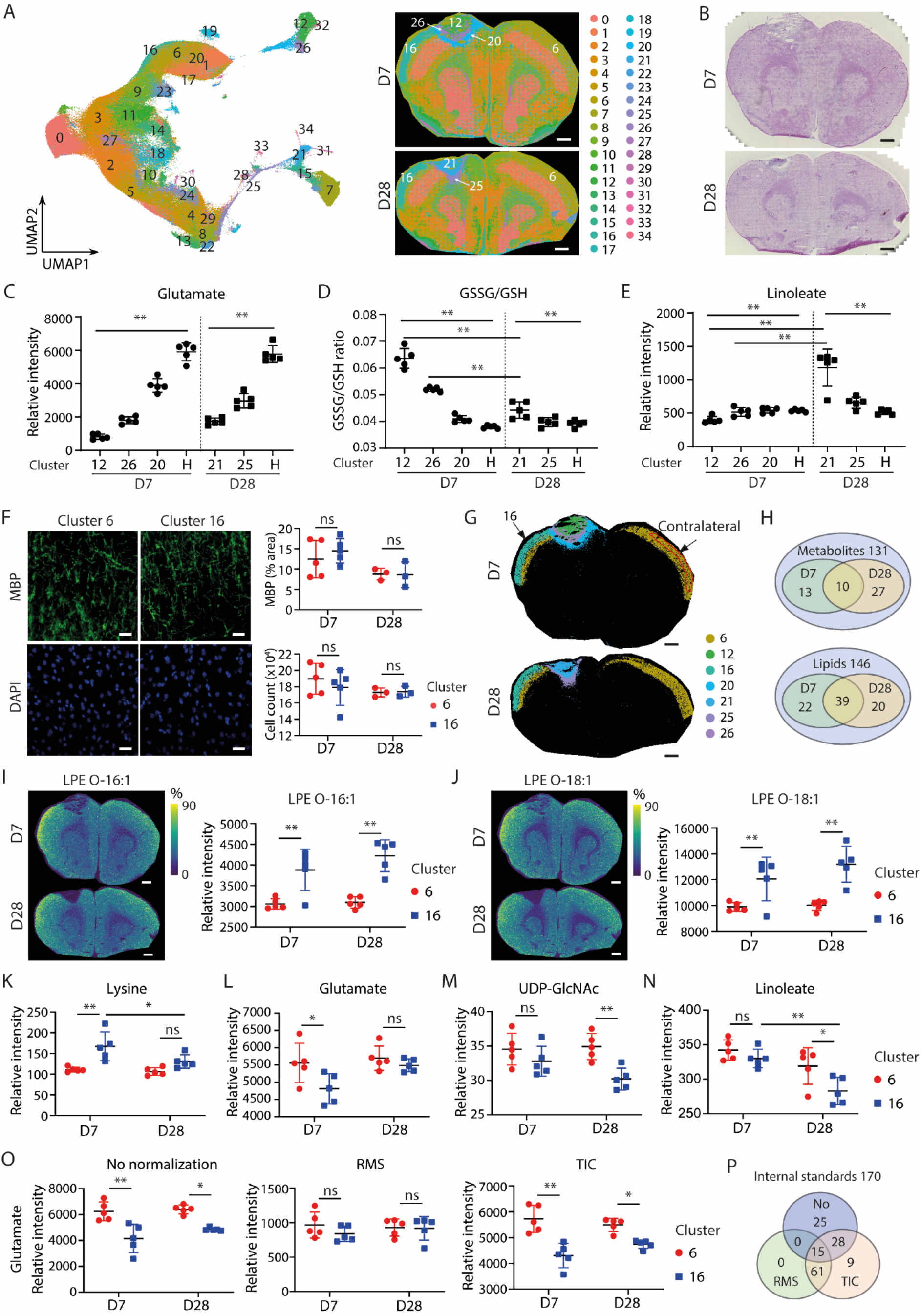
Remote metabolic alterations on ipsilateral cortex after ischemic stroke revealed by quantitative spatial metabolomics. **A,** Brain regional segmentation based on MSI derived lipid profiles of day 7 post-stroke (D7) and day 28 post-stroke (D28) samples (n=5 per group). **B,** H&E staining of post-MSI tissue. **C,** Graphs showing relative quantification of glutamate in infarct (clusters 12, 26 and 21), peri-infarct (clusters 20 and 25) and healthy (H) areas. **D,** Graph showing the ratio of GSSG/GSH in infarct (clusters 12, 26 and 21), peri-infarct (clusters 20 and 25) and healthy (H) areas. **E,** Graph showing relative quantification of linoleate in infarct (clusters 12, 26 and 21), peri-infarct (clusters 20 and 25) and healthy (H) areas. **F,** Representative images of myelin (MBP) and DAPI staining on the cortex areas of D7 samples, scale bar = 20 μm. Graphs showing the comparison of myelin coverage and cell count on different cortex areas of D7 and D28 samples. **G,** Spatial segmentation of the infarct and cortex areas. Cluster 16 and its contralateral region in cluster 6 were highlighted. **H,** Amount of quantified metabolite/lipid species and the significant difference between cluster 16 and its contralateral region within cluster 6. **I,** Representative image showing LPE O-16:1 distribution in the post-stroke brains. Relative quantification in clusters 16 and its contralateral region within cluster 6. **J,** Representative image showing LPE O-18:1 distribution in the post-stroke brains. Relative quantification in clusters 16 and its contralateral region within cluster 6. **K-N,** Relative quantification of lysine **(K)**, glutamate **(L)**, UDP-GlcNAc **(M)** and linoleate **(N)** in cluster 16 and its contralateral region within cluster 6. **O**, Relative intensity of glutamate in cluster 16 and its contralateral region within cluster 6 using different normalization methods. **P**, Amount of internal standards and the significant difference between cluster 16 and its contralateral region within cluster 6 using different normalization methods. All scale bars = 600 μm. *p < 0.05, **p < 0.01.

To validate the superiority of our novel ^13^C-labeled IS based methodology, we also compared these results with the traditional RMS and TIC normalization methods. Comparing the average peak intensities of 4 metabolites of interest in cluster 16 to its contralateral region revealed various different outcomes, of which none was consistent with the IS normalization method (Figure 2O, Supplementary Figures 4A-C), although ion suppression or matrix effect are generally recognized as region specific. This phenomenon indicates that alteration of analytes composition upon injury will affect the ion suppression or matrix effect of MSI measurements even when comparing similar regions. To prove this, we further compared the peak intensities of all 170 homogenously deposited IS between cluster 16 and its contralateral region using different normalization methods. A significant number of IS resulted in altered peak intensity between contralateral regions (Figure 2P, Supplementary Figures 4D-F). Moreover, it is evident that the outcome can be altered depending on the choice of normalization method (Figure 2P). These data suggest an urge for the use of quantitative methods when analyzing possible differences in both inter- and intra-regional tissue sections.

In summary, we introduce a quantitative spatial metabolomics approach utilizing ^13^C-labeled yeast extract as a source of isotopically labeled IS. This method enables both, relative quantification with pixelwise IS normalization and absolute spatial quantification. Beyond metabolite IS, the inclusion of IS for lipids significantly enhances the quantification capacity of this method by orders of magnitude^8, 22^. Using an experimental stroke model we demonstrate that our advanced quantitative methodology does not only surpass classic histological methods but also traditional MALDI-MSI approaches revealing novel, remote regions in the brain that have hitherto evaded histological and molecular imaging. Ion suppression or matrix effects are affected by physico-chemical properties not only in different regions, but also different (patho-) physiological conditions in similar regions. Thus, the choice of different normalization methods leads to altered outcomes, which is also true for the choice of matrix and MSI technique. For example, MALDI-MSI detection of a homogeneously deposited lipid standard (15:0–18:1 (d7) PE) on brain showed distinct MSI images with un-equal distributed peak intensity when comparing norharman or DHB as a matrix^8^. Furthermore, MALDI-MSI and DESI-MSI showed opposite pool-size results of long chain fatty acid palmitate or stearate when comparing tumor relative to healthy tissue regions in a mouse model of glioblastoma^23^. While we demonstrate our method using NEDC matrix and MALDI-MSI, alternative matrices or other analytical modalities such as DESI-MSI should equally benefit from the presented approach. In essence, this method sets a new standard for spatial metabolomics studies, offering improved accuracy and reduced batch effects.

## Methods

### Mouse studies

Mouse kidneys were obtained from normal 12-week-old male DBA/2J mice (n = 3). Animal experiments were approved by the Ethical Committee on Animal Care and Experimentation at the Leiden University Medical Center (Permit No. AVD1160020172926). Control mouse brains were obtained from normal 12-week-old male C57BL/6J mice (n = 3). Animal experiments were approved by the Ethical Committee on Animal Care and Experimentation at the Leiden University Medical Center (Permit No. AVD11600202316801). Mice were perfused with cold PBS-heparin (5IU/mL) via the left ventricle at a controlled pressure of 150 mmHg for 6 minutes to exsanguinate the kidneys or brains before removal. All the tissues were quenched with liquid N2 and stored at -80 °C. All work with animals was performed in compliance with the Dutch government’s guidelines. Mice were housed at 20–22 °C in individually ventilated cages, humidity controlled (55%) with free access to food and water and a light/dark cycle of daytime (06:30–18:00) and nighttime (18.00–06:30).

For Brain stroke experiments, all animal procedures were performed in accordance with the guidelines for the use of experimental animals and were approved by the respective governmental committees (Regierungspraessidium Oberbayern, the Rhineland Palatinate Landesuntersuchungsamt Koblenz). Wild-type C57BL6/J mice were purchased from Charles River. Wild-type male mice (aged 12-13 weeks) were used for the experiments. All mice had free access to food and water at a 12 h dark-light cycle and housed under a controlled temperature (22 +/-2°C). All animal experiments were performed and reported in accordance with the ARRIVE guidelines.

### Photothrombosis experimental stroke surgery

For photothrombosis (PT) induction, mice were anaesthetized with isoflurane, delivered in a mixture of 30% O2 and 70% N2O. Mice were then placed in a stereotactic frame and body temperature was maintained at 37°C with a warming pad. Dexpanthenol eye ointment was applied to both eyes. Animals received 10 μl/g body weight of 1% Rose Bengal (198250-5g; Sigma-Aldrich) in saline i.p. 5 min before the induction of anesthesia (5% isoflurane). A skin incision was made to expose the skull. Bregma was located and the laser collimator was set at 1mm diameter and placed over the lesion location (1.5 mm lateral and 1.0 mm rostral to bregma). 10 min after Rose Bengal injection the laser (25 mV output) was applied to the lesion area for 17 min (Cobolt Jive 50, 561 nm power; Fiber Collimation Package: 543 nm).

Mice were terminally anesthetized with ektamine (120mg/kg) and xylazine (16mg/kg). Mice were then transcardially perfused with 0.9% NaCl and the brain was carefully excised and immediately frozen on dry ice. Brain cryostat sections were cut at 10 μm for Q-MSI and 20 μm for immunohistochemistry and directly mounted on glass slides and stored at -80°C.

### Immunohistochemistry

For immunohistochemical analysis of MBP and Dapi, tissue was fixed on slides with 4% PFA for 20 min at RT, followed by three washes of 1X PBS. Tissue was permeabilized and blocked for 1hr at RT in 1X PBS containing 2% goat serum (16210-064, Thermo Fischer), 2% BSA (A3912, Sigma-Aldrich) and 0.2% Triton X-100 (X-100, Sigma-Aldrich). Sections were then immunolabeled with MBP (1:200, MAB386, Biorad) for 24h at 4°C. After primary antibody labeling, sections were washed three times in 1X PBS for 10 min at RT and incubated with fluorescent secondary antibody (Alexa Fluor 488 goat anti-rat, A11006, Invitrogen) for 2h RT. Sections were washed three times in 1X PBS for 10 min at RT and incubated with Dapi (D1306, Invitrogen) for 5 min at RT. Then washed three more times in 1X PBS for 10 min at RT and sealed with coverslips using Fluoromount medium (F4680, Sigma-Aldrich).

### Confocal microscopy acquisition and image analysis

All images were taken on a Zeiss LSM 880 confocal imaging system with 40x oil immersion objective. Z-stacks were taken at 1 μm intervals at three regions along layers II/III of the cortex per hemisphere. Coverage analysis was performed using the “Particle Analysis” function in Image J. The % area of three images per hemisphere were then averaged.

To quantify cell count, Dapi was used to manually count the nuclei in Image J using the “Multi-point” function. The cell count for the three regions imaged per hemisphere were then averaged.

### Internal Standard and Matrix Application

Tissues were embedded in 10% gelatin and cryosectioned into 10 μm thick sections using a Cryostar NX70 cryostat (Thermo Fisher Scientific, MA, USA) at -20 °C. The sections were thaw-mounted onto indium-tin-oxide (ITO)-coated glass slides (VisionTek Systems Ltd., Chester, UK). Mounted sections were placed in a vacuum freeze-dryer for 15 minutes prior to matrix application. After drying, the tissue slices on ITO slides were heated to ∼80 °C for 15 s on a hot plate to denature metabolic enzymes.

The unlabeled metabolite yeast extracts (Cambridge Isotope Laboratories, Inc. ISO1-UNL) and U-^13^C-labeled metabolite yeast extracts (99%, Cambridge Isotope Laboratories, Inc. ISO1) were reconstituted in 2 mL 50% methanol/deionized water. The mixtures were shaken by hand with intermittent high-speed vortexing for a minimum of 2 min, and followed by centrifuge at 20 °C for 5 min at 4000 rcf. The clear standard solutions were collected and stored at -80 °C.

For IS applications, the ^13^C-labeled yeast extract solution was diluted in methanol (1/10 v/v). Then, IS were sprayed on the tissue surface using a HTX M3+ Sprayer (HTX Technologies, USA). After application of IS, N-(1-naphthyl) ethylenediamine dihydrochloride (NEDC) (Sigma-Aldrich, UK) MALDI-matrix solution of 7 mg/mL in methanol/acetonitrile/deionized water (70, 25, 5 %v/v/v) was applied using a HTX M3+ Sprayer (HTX Technologies, USA).

### MALDI-MSI measurement

MALDI-MSI was performed using a TimsTOF MALDI2 system (Bruker Daltonics GmbH, Bremen, Germany). The instrument was calibrated using red phosphorus prior to each measurement. All the data were acquired in negative mode with MALDI2 at a mass range of *m/z* 50–1000. The MALDI2 laser was with a pulse delay time of 10 μs. For kidney and brain stroke tissues, data were acquired at a pixel size of 20 μm (x, y) using a beam scan area of 16 × 16 μm and laser was operated at 1 kHz with 200 laser shots accumulated per pixel. For absolute quantitative analysis on brain tissues, data were acquired at a pixel size of 30 μm (x, y) using a beam scan area of 26 × 26 μm and laser was operated at 1 kHz with 100 laser shots accumulated per pixel.

### Quantification of metabolites from unlabeled yeast extract by NMR

An aliquot of unlabeled yeast extract was prepared in 0.05 M phosphate buffer in 99.8% deuterated water including 0.05mM of trimethylsilyl propionic-d4-sodium salt (TSP-d4) as reference and quantification standard. 185 μL was transferred to a 3 mm NMR tube and one NMR experiment (pulse sequence: noesygppr1d; Bruker Biospin Ltd) was collected in a 14.1 T (600 MHz for 1H) Bruker Avance Neo NMR. Yeast metabolites were then quantified using the Chenomx NMR suite 10.0 software (Chenomx NMR suite, v10.0, Edmonton, AB, Canada). IDs of metabolites were confirmed by 2D NMR (1H-1H TOCSY and 1H-13C HSQC) experiments.

### Untargeted LC-MS/MS metabolomics measurements

Normal kidney and brain samples stored at -80 °C were thawed on ice. The thawed sample was homogenized in a grinder (30 Hz) for 20 s. A 400 μL solution (methanol : water = 7:3, v/v) containing IS was added to 20 mg homogenized tissue, and shaken at 1500 rpm for 5 min. After placing on ice for 15 min, the sample was centrifuged at 12000 rpm for 10 min (4 °C). A 300 μL aliquot of supernatant was collected and placed at -20 °C for 30 min. The sample was again centrifuged at 12000 rpm for 3 min (4 °C). Finally, a 200 μL aliquot of supernatant was transferred for LC-MS analysis.

Unlabeled yeast extract stored at -80 °C was thawed on ice and vortexed for 10 s. A 150 μL extract solution (acetonitrile : nmethanol = 1:4, v/v) containing IS was added to 50 μL sample. The sample was vortexed for 3 min and centrifuged at 12000 rpm for 10 min (4 °C). A 150 μL aliquot of the supernatant was collected and placed at -20 °C for 30 min, and again centrifuged at 12000 rpm for 3 min (4 °C). Finally a 120 μL aliquot of supernatant was transferred for LC-MS analysis.

Samples were analyzed with two LC/MS methods. One aliquot was analyzed using positive mode electrospray ionization employing aT3 column (Waters ACQUITY Premier HSS T3 Column 1.8 μm, 2.1 mm × 100 mm) using 0.1 % formic acid in water as solvent A and 0.1 % formic acid in acetonitrile as solvent B running the following gradient: 5 to 20 % in 2 min, increased to 60 % in the following 3 min, increased to 99 % in 1 min and held for 1.5 min, return to 5 % mobile phase B within 0.1 min, held for 2.4 min. The column oven was kept at 40 °C; the flow rate was 0.4 mL/min; the injection volume was 4 μL. Using identical conditions a second aliquot was analyzed in negative mode electrospray ionization mode.

The mass spectrometer operated under the following conditions, data acquisition was operated using the information-dependent acquisition (IDA) mode using Analyst TF 1.7.1 Software (Sciex, Concord, ON, Canada). The source parameters were set as follows: ion source gas 1 (GAS1) 50 psi; ion source gas 2 (GAS2) 50 psi; curtain gas (CUR) 25 psi; temperature(TEM) 550 °C; declustering potential (DP) 60 V, or−60 V in positive or negative mode, respectively; and ion spray voltage floating (ISVF), 5000 V or−4000 V in positive or negative modes, respectively. The TOF MS scan parameters were set as follows: mass range, 50–1000 Da; accumulation time, 200 ms; and dynamic background subtract, on. The product ion scan parameters were set as follows; mass range, 25–1000 Da; accumulation time, 40 ms; collision energy, 30 or−30 V in positive or negative modes, respectively; collision energy spread, 15; resolution, UNIT; charge state, 1 to 1; intensity, 100 cps; exclude isotopes within 4 Da; mass tolerance, 50 ppm; maximum number of candidate ions to monitor per cycle, 18.

### Untargeted LC-MS/MS lipidomics measurements

Normal kidney and brain samples stored at -80 °C were thawed on ice. The thawed 20 mg of sample was homogenized in a grinder (30 Hz) for 20 s and the centrifuge (3000 rpm, 4°C) for 30 s. For unlabeled yeast extract stored at -80 °C was thawed on ice and vortexed for 10 s. Then mix the tissue extract or 200 μL of the yeast extract with 1mL of the extraction solvent (tert-butyl methyl ether: methanol = 3:1, v/v) containing IS mixture. After whirling the mixture for 15 min, 200 μL of ultrapure water was added. Vortex for 1 min and centrifuge at 12,000 rpm for 10 min. 200 μL of the upper organic layer was collected and evaporated using a vacuum concentrator. The dry extract was dissolved in 200 μL reconstituted solution (acetonitrile: isopropanol=1:1, v/v) to LC-MS/MS analysis.

The sample extracts were analyzed using an LC-ESI-MS/MS system (UPLC, ExionLC AD; MS, QTRAP® 6500+ System). The analytical conditions were as follows, UPLC: column, Thermo Accucore™C30 (2.6 μm, 2.1 mm×100 mm i.d.); solvent system, A: acetonitrile/water (60/40,V/V, 0.1% formic acid, 10 mmol/L ammonium formate), B: acetonitrile/isopropanol (10/90 VV/V, 0.1% formic acid, 10 mmol/L ammonium formate); gradient program, A/B (80:20, V/V) at 0 min, 70:30 V/V at 2.0 min, 40:60 V/V at 4 min, 15:85 V/V at 9 min, 10:90 V/V at 14 min, 5:95 V/V at 15.5 min, 5:95 V/V at 17.3 min, 80:20 V/V at 17.3 min, 80:20 V/V at 20 min; flow rate, 0.35 ml/min; temperature, 45 °C; injection volume: 2 μL. The effluent was alternatively connected to an ESI-triple quadrupole-linear ion trap (QTRAP)-MS.

LIT and triple quadrupole (QQQ) scans were acquired on a triple quadrupole-linear ion trap mass spectrometer (QTRAP), QTRAP® 6500+ LC-MS/MS System, equipped with an ESI Turbo Ion-Spray interface, operating in positive and negative ion mode and controlled by Analyst 1.6.3 software (Sciex). The ESI source operation parameters were as follows: ion source, turbo spray; source temperature 500 °C; ion spray voltage (IS) 5500 V (Positive), -4500 V (Neagtive); Ion source gas 1 (GS1), gas 2 (GS2), curtain gas (CUR) were set at 45, 55, and 35 psi, respectively. Instrument tuning and mass calibration were performed with 10 and 100 μmol/L polypropylene glycol solutions in QQQ and LIT modes, respectively. QQQ scans were acquired as MRM experiments with collision gas (nitrogen) set to 5 psi. DP and CE for individual MRM transitions was done with further DP and CE optimization. A specific set of MRM transitions were monitored for each period according to the metabolites eluted within this period.

### MSI data processing and analysis

MSI data were exported and processed in SCiLS Lab 2023c Pro (SCiLS, Bruker Daltonics). Peak picking was performed on the average spectrum with a threshold signal-to-noise-ratio > 3 and relative intensity > 0.01% in mMass, and matrix peaks were excluded from the *m/z* feature list. The metabolite *m/z* values from unlabeled yeast extract after peak picking were compared with the untargeted LC-MS/MS dataset after re-calibration in mMass and annotated with an error < ±10 ppm. The *m/z* features, which were not found in the LC-MS/MS dataset, were further imported into the Yeast Metabolome Database and Human Metabolome Database for potential annotation with an error < ±10 ppm. The lipid *m/z* values from unlabeled yeast extract were imported into the LIPID MAPS database and preliminarily annotated with an error < ±10 ppm. Odd-chain lipids were removed as they are very rare in mammals. Most of the annotated metabolites and lipids were further confirmed with the certificate of the product from the supplier. For kidneys and brain tissues, peaks overlapping with ^13^C-labeled yeast extract, but not with the control tissue, were removed. In summary, selected *m/z* values were annotated by the following strategy: i) database search using high resolution masses against the Human Metabolome Database, and LIPID MAPS database with an error < ±10 ppm, ii) validation of the so generated hits by off-tissue LC-MS/MS analysis and iii) comparison with a previously published dataset^24^ to further verify the annotations..

After annotation of the *m/z* features from un-labeled yeast extracts, theoretical *m/z* values of the uniformly ^13^C-labeled metabolites and lipids were calculated based on adduct *m/z* values and chemical formula of the annotated metabolites and lipids. For U-^13^C-labeled yeast extract, the peaks overlapping with peaks of un-labeled yeast extracts were removed during peak picking. The *m/z* values of U-^13^C-labeled yeast extract after peak picking were further compared with the calculated theoretical *m/z* value of the uniformly ^13^C-labeled metabolites and lipids. The uniformly ^13^C-labeled metabolites and lipids found in ^13^C-labeled yeast extract were annotated with an error < ±10 ppm. A library of IS was established containing un-labeled metabolites and their uniformly ^13^C-labeled counter parts.

Subsequently this library was applied for the analysis of MSI data from kidneys and brains. Peaks from kidney or brain tissues matching the so established library were selected for further analysis. Subsequent quality control included a final removal of overlapping ^13^C labelled signals with endogenous non-labelled signals. Finally, the remaining IS used for pixelwise normalization were selected on both kidney and brain tissues. The metabolite IS were used for relative quantification of their paired un-labeled metabolites on tissues. The selected lipid IS with highest peak intensity were used for relative quantification of 8 different lipid classes. The pixelwise normalization to IS was performed using in SCiLS Lab 2023c Pro. Peak intensities after normalization of the selected features were exported for all the measured pixels from SCiLS Lab 2016b for the following analysis.

For absolute quantitative spatial metabolomics analysis, eight different amounts of unlabeled yeast extracts were spotted next to the tissue sections with two replicates for each concentration. The absolute amount of the metabolites on each spot was calculated based on NMR measurement. The peak intensity of unlabeled metabolite from spotted unlabeled yeast extract was normalized to its paired ^13^C-labled IS, subsequently the normalized intensity was exported for all measured pixels in SCiLS Lab 2023c Pro. The normalized peak intensity and its corresponding concentration were used to generate the calibration curves with individual equations for each metabolites in GraphPad Prism 9. The equations were further used for absolute quantification in each pixel from the tissues.

The *m/z* features of endogenous lipids or *m/z* features of exogenous metabolites and lipids were selected for UMAP analysis. Their intensities were exported for all the measured pixels with root mean square normalization from SCiLS Lab 2016b. The datasets were transformed into a count matrix by taking the integer. This count data matrix was normalized and scaled using SCTransform to generate a 2-dimensional UMAP map using Seurat in R. The distribution of the pixels from different clusters on tissues were co-registered to the post-MSI staining, and regions were identified based on both staining and their morphology. For the stroke study, the region information of each pixel was used to calculate the average abundance of both the endogenous metabolites and IS for regional comparison. For cluster 16, its contralateral regions within cluster 6 were selected manually in SCiLS Lab 2023c Pro after co-registering with the cluster distribution (Figure 2G). The average abundance of both metabolites and IS from cluster 16 and its contralateral region were exported from SCiLS Lab 2023c Pro for further comparison.

For molecular histology, the datasets were used to generate a 3-dimensional UMAP map using packages Seurat 3.0 and plotly. The embedding information of the 3-dimensional UMAP was translated to RGB color coding by varying red, green and blue intensities on the 3 independent axes. Together with pixel coordinate information exported from SCiLS Lab 2023c Pro, a MxNx3 data matrix was generated and used to generate UMAP images in Matlab R2019a.

### Statistics and reproducibility

All the experiments and data analysis were performed on 3 or 5 animals per group. No animals or data points were excluded from the analysis. All the data are presented as mean ± SD, unless indicated otherwise. Data normality and equal variances were tested using Shapiro-Wilk test.

Differences between over three groups were assessed by one way ANOVA test following by TukeyHSD test. Differences between two groups were assessed by 2-tailed Student’s t test, when not normally distributed, by two-tailed F-test. P-value < 0.05 were considered statistically significant.

## Acknowledgements

TR is supported by the Novo Nordisk Foundation Center for Stem Cell Medicine (reNEW, supported by Novo Nordisk Foundation grant (NNF21CC0073729)) and is funded by the European Union. Views and opinions expressed are however those of the author(s) only and do not necessarily reflect those of the European Union or the European Research Council Executive Agency. Neither the European Union nor the granting authority can be held responsible for them. This work is supported by ERC grant (SPARK 101140863).

## Author Contributions

GW designed research study, conducted experiments, processed data, wrote the manuscript. BMvdB performed the animal experiments and gave comments. SK performed the NMR experiments and gave comments. KP and AL provided the mouse stroke samples, performed the IF staining and gave comments. MJ performed the post-MSI staining and gave comments. MG designed research study and wrote the manuscript. TJR organized funding, designed research study and wrote the manuscript.

---

**Supplementary figure 1.**
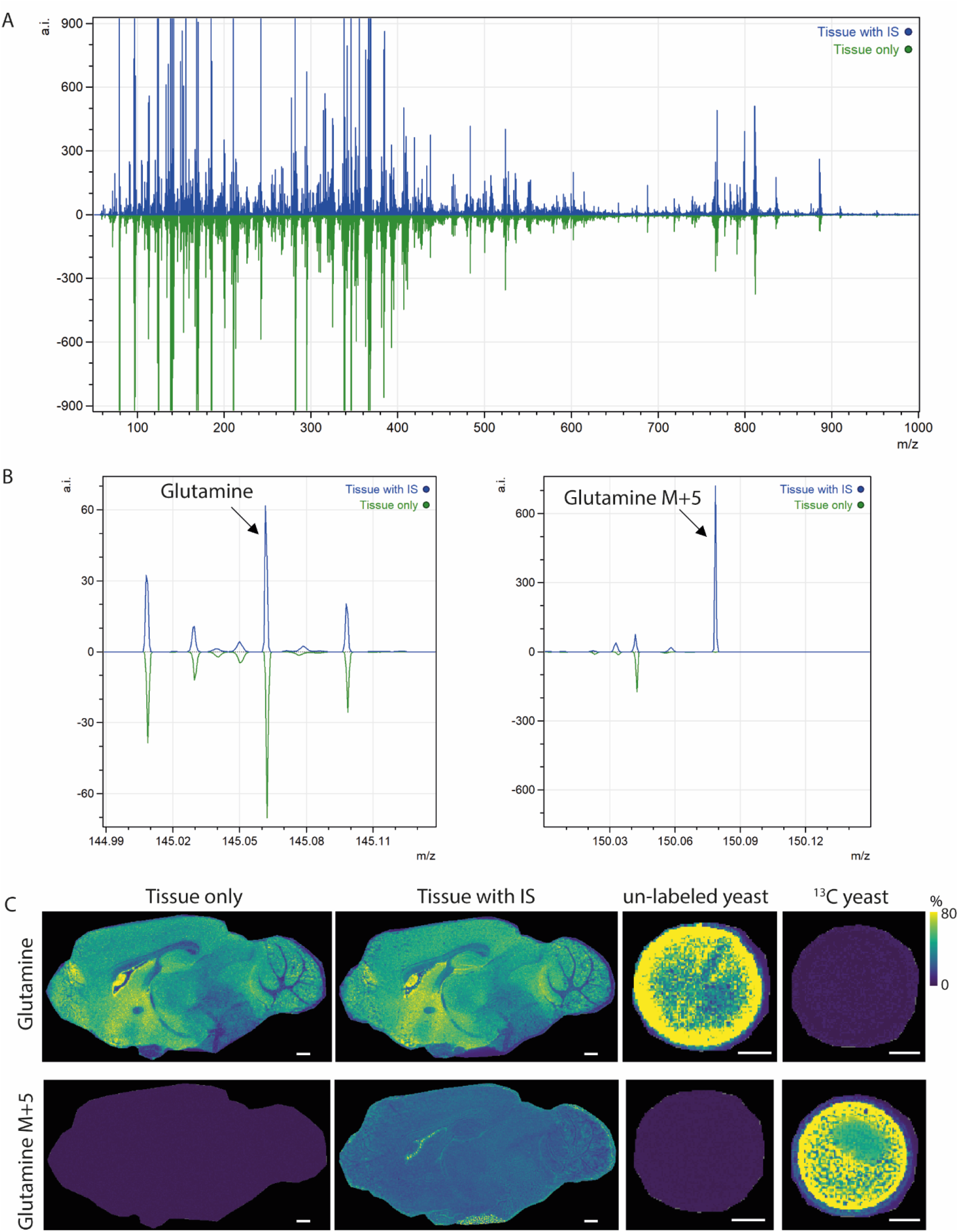
Spectrum for ^13^C yeast-assisted mass spectrometry. **A,** Comparison of spectra from kidney tissues sprayed with and without ^13^C-labeled yeast extract. **B,** Glutamine peaks from spectra of kidney tissue sprayed with and without ^13^C-labeled yeast extract. **C,** Comparison of glutamine distribution from brain tissues sprayed with and without ^13^C-labeled yeast extract and spotted different yeast extracts. All scale bars = 600 μm.

**Supplementary figure 2.**
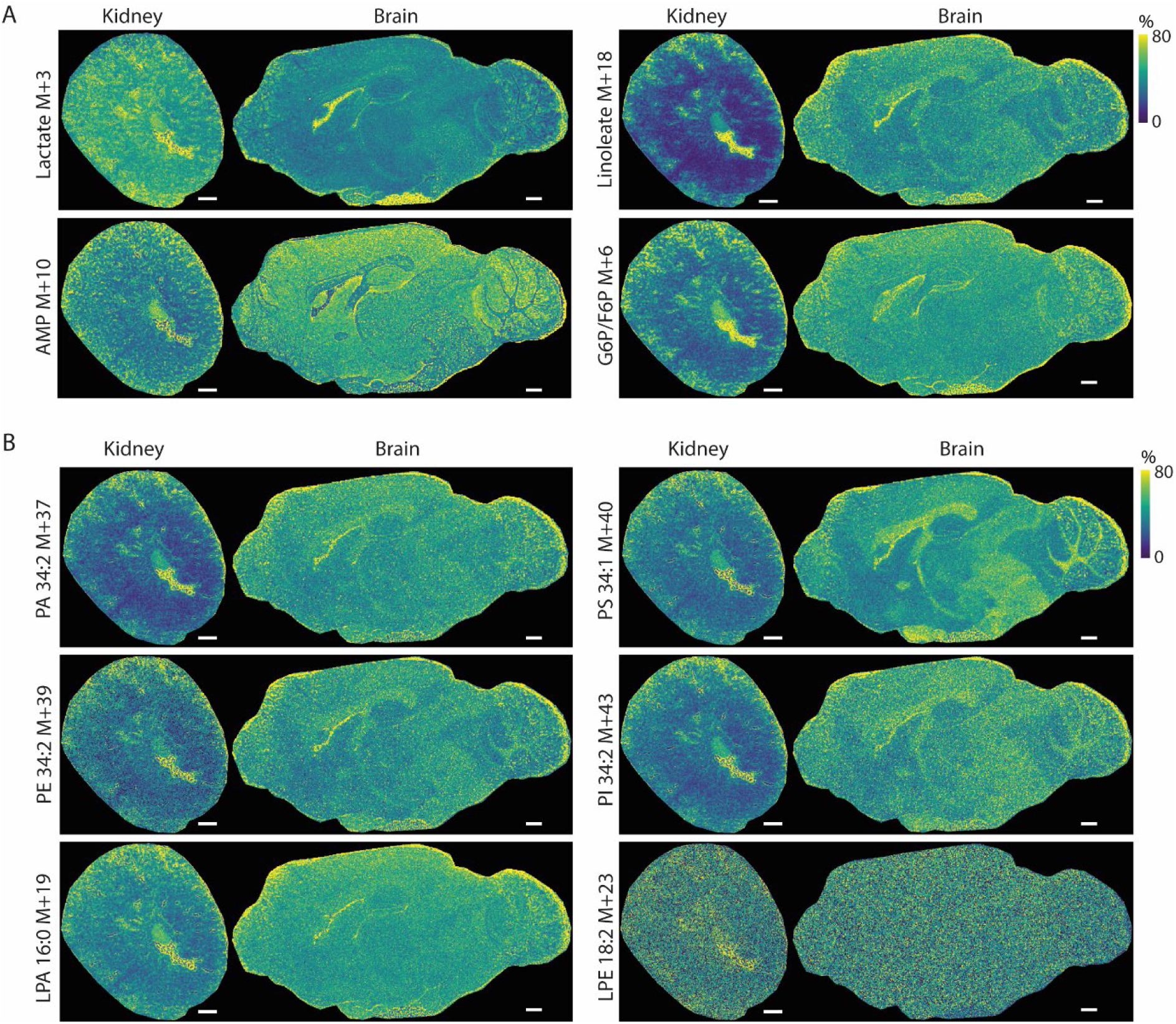
Ion suppression of different metabolites on both kidney and brain. **A,** Distribution of different metabolite internal standards from ^13^C-labeled yeast extract sprayed on both kidney and brain tissues. **B,** Distribution of internal standards representing 6 lipids classes from ^13^C-labeled yeast extracts sprayed on both kidney and brain tissues. All scale bars = 600 μm.

**Supplementary figure 3.**
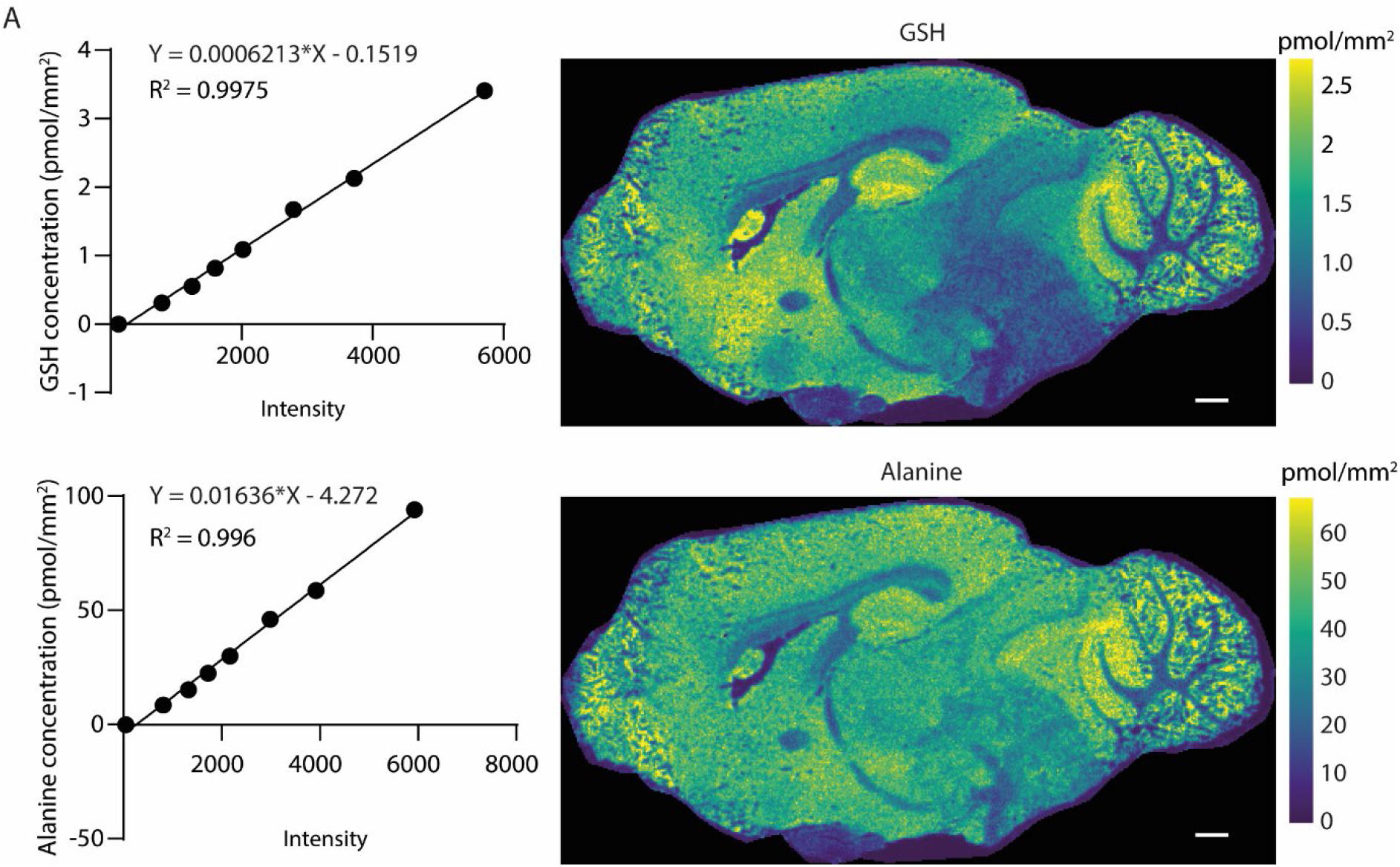
Absolute spatial quantification of metabolites on mouse brain. **A,** Calibration curve and absolute spatial quantification of glutathione (GSH) and alanine on mouse brain.

**Supplementary figure 4.**
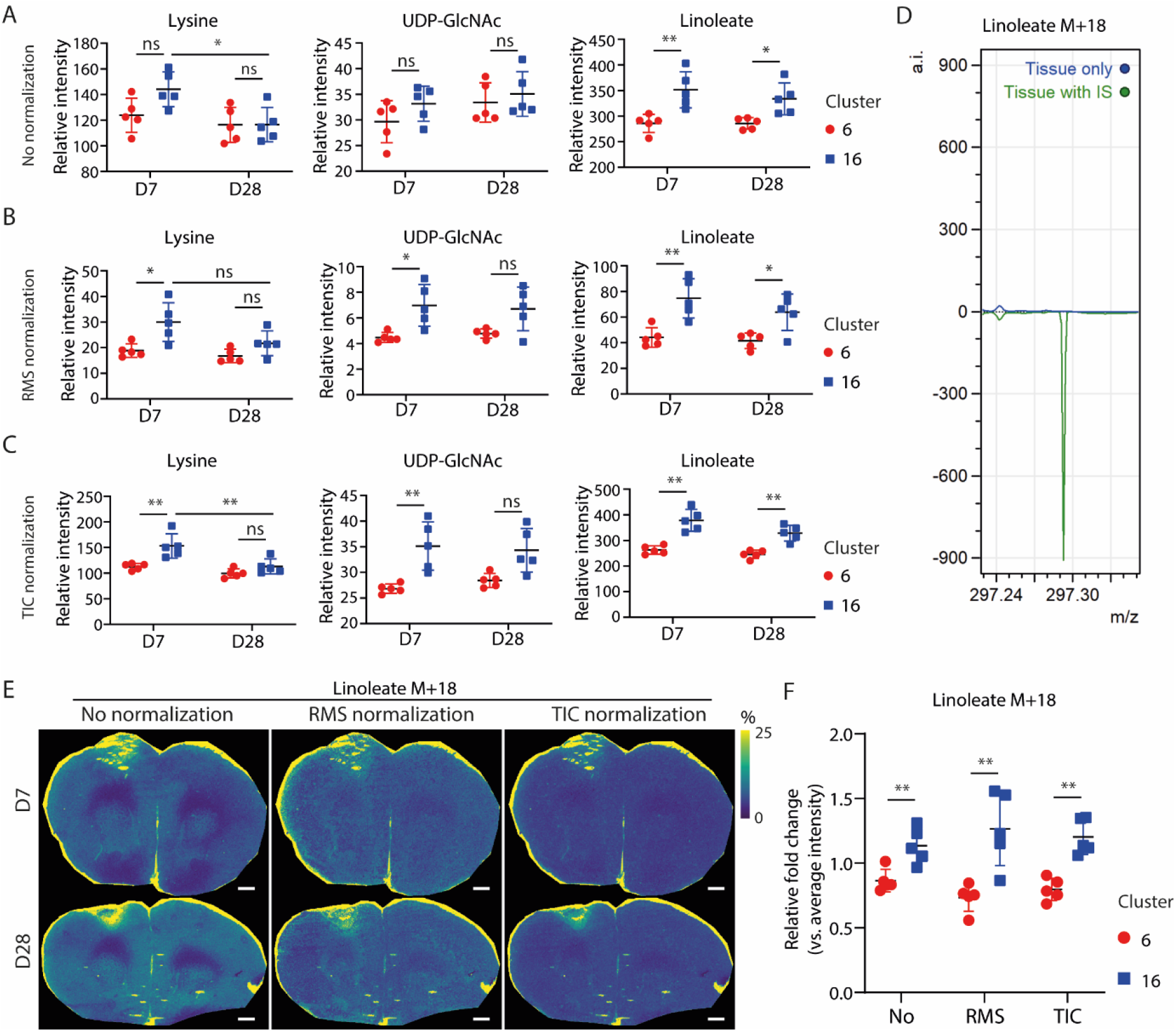
comparison of three traditional normalization methods. **A,** Relative peak intensity of different metabolites in clusters 16 and its contralateral region within cluster 6 without normalization. **B**, Relative peak intensity of different metabolites in clusters 16 and its contralateral region within cluster 6 using RMS normalization. **C**, Relative peak intensity of different metabolites in clusters 16 and its contralateral region within cluster 6 using TIC normalization. **D**, Comparison of linoleate M+18 spectra from brain stroke sample sprayed with and without ^13^C-labeled yeast extract. **E**, Comparison of linoleate M+18 distribution on brain stroke samples sprayed with 13C-labeled yeast extract using different normalization methods. **F**, Comparison of linoleate M+18 relative intensity in clusters 16 and its contralateral region using different normalization methods. All scale bars = 600 μm. *p < 0.05, **p < 0.01.

